# 5-methylcytosine and 5-hydroxymethylcytosine are synergistic biomarkers for early detection of colorectal cancer

**DOI:** 10.1101/2024.10.30.621123

**Authors:** Fabio Puddu, Annelie Johansson, Aurélie Modat, Jamie Scotcher, Riccha Sethi, Shirong Yu, Nick Harding, Mark Hill, Ermira Lleshi, Casper Lumby, Jean Teyssandier, Michael Wilson, Robert Crawford, Tom Charlesworth, Robert J Osborne, Shankar Balasubramanian, Páidí Creed

## Abstract

Early cancer detection has the potential to significantly improve treatment outcomes and survival rates. This study investigates the roles of 5-methylcytosine (5mC) and 5-hydroxymethylcytosine (5hmC) as biomarkers for early-stage colorectal cancer (CRC) detection in cell-free DNA (cfDNA). Using whole genome sequencing, we analyzed cfDNA from 37 treatment-naive CRC patients and 32 healthy controls. Our findings indicate that combining measurements of 5mC and 5hmC significantly enhances diagnostic accuracy (AUC = 0.95) compared to traditional approaches that conflate these markers (modified C, AUC = 0.66). Notably, 71.7% of differentially methylated regions (DMRs) exhibiting an increase in 5hmC in stage I cfDNA also showed a corresponding decrease in 5mC in stage IV, suggesting that 5hmC can effectively track regions undergoing demethylation during tumor development. These results support the hypothesis that distinguishing between 5mC and 5hmC can improve the sensitivity of liquid biopsy tests for early cancer detection.

## Introduction

Early detection has the potential to transform the treatment and survival of cancer (Crosby et al, 2022). Cancer is caused by changes to the genome and epigenome, characterized by somatic mutations (Stratton et al., 2009, Lawrence et al., 2014) and epigenetic reprogramming (Brock et al., 2009; Timp and Feinberg., 2013; Flavahan et al., 2017; Marine et al., 2020). Cancer-associated somatic mutations affect epigenetic systems in hematological and solid tumors (Hutter and Zenklusen, 2018; CGAR, 2013; Feinberg et al., 2016), and enzymes that alter epigenetic modifications are targeted by epi-drugs in cancer therapy (Bates et al., 2020). Recent evidence also suggests that epigenetic perturbation can drive cancer initiation independently of somatic mutations (Parreno et al, 2024).

DNA cytosine methylation and demethylation result in chemically stable epigenetic changes that can be accurately measured. These modifications predominantly occur at CpG dinucleotides, where cytosines are followed by guanines in the DNA sequence. Cytosine methylation results in 5-methylcytosine (5mC) and enzymatic oxidation converts 5mC to 5-hydroxymethylcytosine (5hmC), which upon further oxidation generates cytosine derivatives that are removed by repair pathways resulting in overall demethylation (Kriaucionis and Heintz, 2009; Tahiliani et al., 2009; He et al., 2011; Ito et al., 2011) (**Figure 1a**). 5mC and 5hmC play distinct roles in epigenetic regulation, although this information is often lost as they are (mis)read as a combined signal in classical sequencing approaches. 5mC is typically found at gene promoters where it is associated with gene repression (Jones, 2012). In contrast, 5hmC is enriched at poised or active enhancers and gene bodies of actively transcribed genes and plays a critical role in the transcriptional reprogramming of cells that are undergoing state transitions (Stroud et al., 2011; Hahn et al., 2013).

**Figure 1.**
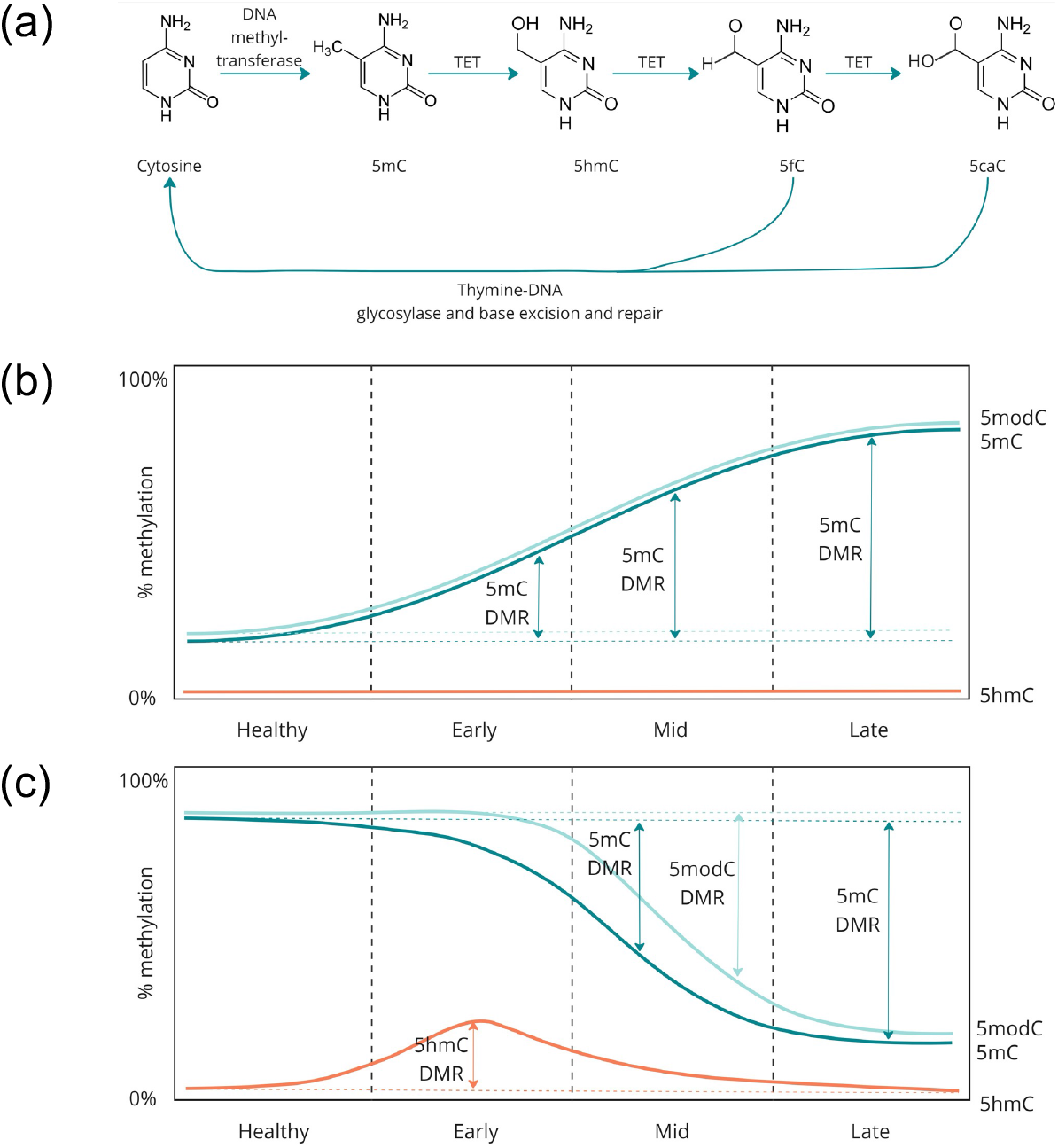
(a) Schematic of the enzymatic steps involved in cytosine methylation and demethylation. DNA methyltransferases catalyse the addition of a methyl group to a cytosine base in a CpG context. The methyl group is oxidised by TET enzymes to 5hmC, and downstream oxidised derivatives 5fC and 5caC, before thymine DNA glycosylase mediates the removal of 5fC or 5caC, with subsequent replacement with unmodified cytosine by gap-filling repair. (b) Schematic representation of the increase in 5mC leading to hypermethylation or (c) loss of 5mC leading to hypomethylation in a region of the genome. (b) Represents a region with 5mC hypermethylation in late-stage cancer, where 5mC accumulates during disease progression. For these regions, changes in 5mC and modC are approximately equivalent and would therefore be expected to have similar power as biomarkers to distinguish between disease stages and healthy individuals. (c) Represents a region with hypomethylation in late-stage disease. Changes in 5hmC are proportionally larger at the early stage than changes in 5mC or modC. At later stages, as demethylation completes, 5mC and modC become proportionally larger. Changes in modC are masked by the conflation of 5mC and 5hmC and only become distinguishable at mid-to late-stage.

Methylation is used as a biomarker for early cancer detection. Epigenetically reprogrammed tumor cells release DNA into the bloodstream, where methylation changes can be detected in cell-free DNA (cfDNA) (Liu et al., 2020; Chung et al., 2024). Despite progress, early-stage cancers remain difficult to detect. For example, Klein et al (2021) reported methylation-based liquid biopsy data for CRC with 99.5% specificity, showing 43.3% sensitivity at stage I compared to 95.3% sensitivity at stage IV (**Table S1**). The observed lower sensitivity for early-stage cancer, compared to later stages, could be due to the limited amount of tumor-derived cfDNA, subtler epigenetic shifts at this stage as measured by modC (where modC is the combined signal from 5mC + 5hmC), or a combination of both factors. This issue is compounded by the limitations of current methods used to assay methylation. For example, enrichment approaches typically resolve DNA fragments, rather than individual bases, and suffer from limited accuracy in identifying and measuring differentially methylated regions (DMRs) (Ficz et al., 2011; Walker et al., 2022). In addition, bisulfite sequencing conflates 5mC and 5hmC into a single modC signal, masking the individual contributions of these two modifications particularly in early-stage disease (Guerin et al, 2024; **Figure 1b,c**).

Given the different biological roles of 5mC and 5hmC, we hypothesized that discriminating these changes at base resolution would provide more informative data and increase the sensitivity of early-cancer detection. To test this hypothesis, we performed whole genome sequencing (WGS) of liquid biopsy samples from stage I CRC cancer patients and healthy controls, using a method that simultaneously sequences 5mC and 5hmC at read level (Füllgrabe et al., 2023). Our results demonstrate that 5hmC provides informational value that is additive to 5mC, with models utilizing both 5mC and 5hmC achieving higher diagnostic accuracy than modC (AUC 0.95 versus 0.66).

## Results

To test our hypothesis that separate measurement of 5mC and 5hmC would provide more useable information to detect the presence of early-stage cancer in plasma, we examined differences in 5mC and 5hmC in cfDNA from individuals with colorectal cancer and healthy controls. For this purpose, we analysed cfDNA from 69 samples including 32 healthy controls and 37 treatment naive CRC patients (26 stage I and 11 stage IV). Demographics of the cohort including age, sex, and relevant clinical characteristics such as smoking history, are outlined in **Table 1**. For each sample, 10 ng of cfDNA was processed using the duet evoC assay (biomodal), and libraries were subjected to WGS to a mean depth of 34x.

**Table 1.** Demographics and covariates in the cohort.

|  | **Total** | **Control** | **CRC**<br>**stage I** | **CRC stage IV** |
| --- | --- | --- | --- | --- |
| N | 69 | 32 | 26 | 11 |
| Age (years), mean (SD) | 60.8 (6.6) | 58.8 (6.9) | 61.5 (6.0) | 65.5 (4.8) |
| Female gender, n (%) | 48 (83) | 25 (78) | 23 (88) | 9 (82) |
| Ethnicity, n (%) |  |  |  |  |
| White | 69 (100) | 32 (100) | 26 (100) | 11 (100) |
| Alcohol consumption, n (%) |  |  |  |  |
| Unknown | 6 (9) | 5 (16) | 1 (4) | 0 (0) |
| None | 17 (25) | 3 (9) | 8 (31) | 6 (55) |
| Light ≤ drink/day | 46 (67) | 24 (75) | 17 (65) | 5 (46) |
| Smoking, n (%) |  |  |  |  |
| Never smoked | 57 (83) | 24 (75) | 23 (89) | 10 (91) |
| Ex-smoker | 6 (9) | 4 (13) | 2 (8) | 0 (0) |
| Smoker | 6 (9) | 4 (13) | 1 (4) | 1 (9) |

To focus our analysis on regions where significant epigenetic shifts occur, we first identified regions that showed significant differences in modC using publicly available data from 9 stage IV CRC samples with matched normal tissue analyzed by Illumina 450K arrays. Our goal was to track regions undergoing methylation changes, which may be associated with tumor progression. Overall, we identified 11,691 modC DMRs (q < 0.05), of which 7,954 were hypo-methylated and 3,737 were hyper-methylated in CRC (**Figure 2a**). Each of these regions contained one or more CpGs and spanned tens to thousands of base pairs (bp), with a mean size of 857 bp (**Figure S1**).

**Figure 2.**
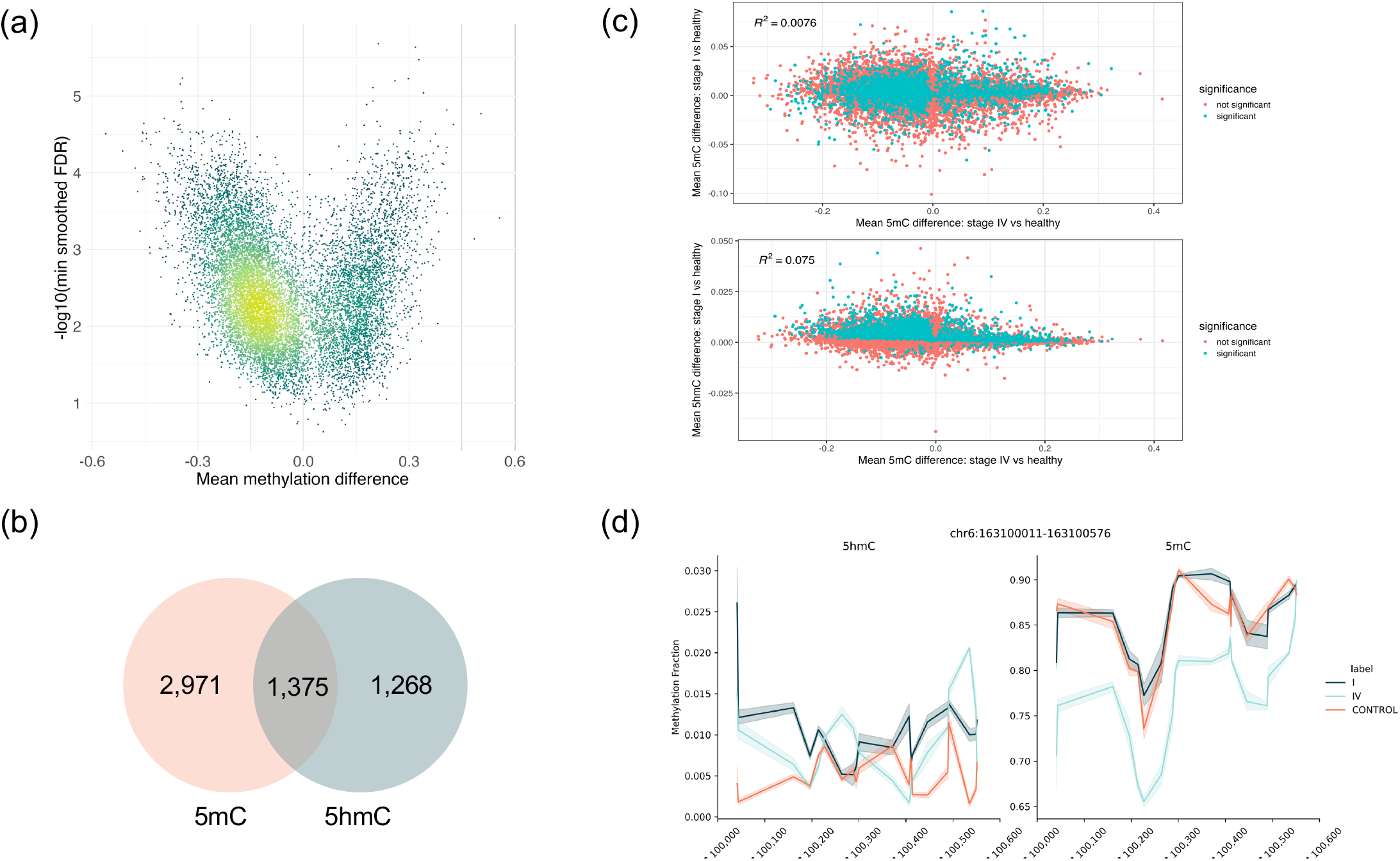
(a) Volcano plot for DMRs between stage IV CRC tissue and adjacent matched normal, using data from TCGA. Colour indicates density of points, with yellow used to indicate a higher density. (b) Venn diagram of regions with statistically significant (q < 0.05) differences in 5mC and 5hmC in stage I plasma. (c) Scatter plot comparing 5mC (upper panel) and 5hmC (lower panel) differences measured in stage I and stage IV plasma. Statistical significance (q < 0.05) at stage I is indicated by blue colour. (d) Trace plots of 5hmC (left) and 5mC (right) fractions in an example region that showed increased 5hmC in stage I plasma and decreased 5mC in stage IV plasma.

We then analyzed differences in 5mC, 5hmC, and modC between plasma from CRC patients and healthy controls across the 11,691 modC DMRs identified in public tissue data, performing separate comparisons for stage I and stage IV plasma. We found a strong correlation between 5mC and modC DMRs in stage IV plasma and the stage IV modC tissue data (both R^2^=0.66). However, no correlation was observed when comparing 5mC or modC DMRs in stage I plasma with stage IV tissue (both R^2^ < 0.001) (**Figure S2**). This difference can be explained by methylation changes during tumor progression, in addition to the higher proportion of tumor DNA in the cfDNA of individuals with late-stage cancer (Diehl et al 2008, Bettegowda et al 2014).

After confirming that 5mC DMRs in stage IV tissue were similarly altered in stage IV plasma, we analyzed 5mC and 5hmC DMRs in stage I plasma to assess their relationship to the changes observed in stage IV plasma. In doing so, we sought to determine whether patterns of changes in 5mC and/or 5hmC at stage I preceded the methylation changes seen in late-stage cancer, potentially acting as an early indicator of ctDNA presence in plasma. In total, 10,557 modC tissue DMRs showed statistically significant differences in 5mC in stage IV plasma (q < 0.05). Of these, we found that 41.2% (4,346/10,557) also had a significant difference in 5mC, and 25% (2,643/10,557) also had a significant difference in 5hmC, in stage I plasma. While there was some overlap between the sets of stage I 5mC and 5hmC DMRs, 48% (1,268/2,643) of the 5hmC DMRs did not show statistically significant differences in 5mC. This suggests that measuring 5hmC in stage I plasma provides complementary information to that obtained from 5mC (**Figure 2b**).

We then compared the direction of DMRs (i.e., increase or decrease in methylation) between stage I and IV plasma. Of the 4,346 5mC DMRs in stage I plasma, 44.6% (1,938/4,436) were consistently methylated or demethylated in both stage I and stage IV plasma. Among the regions which did not show the same direction of change in 5mC in stage I and stage IV, a majority (43.9%; 1,908/4,436) had higher levels of 5mC in stage I plasma and lower levels of 5mC in stage IV plasma (**Figure 2c**).

Given 5hmC is an intermediate of de-methylation (**Figure 1a**), it follows that changes in 5hmC at stage I could precede loss of 5mC at stage IV. Supporting this, the majority (71.7%; 1,894/2,643) of DMRs with statistically significant changes in 5hmC in stage I and 5mC in stage IV plasma had an increase in 5hmC at stage I and a decrease in 5mC at stage IV. Of the remaining 5hmC DMRs, 26% (687/2,643) showed an increase in 5hmC in stage I and an increase in 5mC in stage IV plasma. A small proportion of the regions (2.3%; 62/2,643) showed a statistically significant decrease in 5hmC in stage I plasma. Collectively, these data are consistent with 5hmC being an intermediate in the transition from methylated to unmethylated C during the progression of disease from stage I to stage IV. In particular, a discernible increase in 5hmC in stage I plasma in these regions appears to be a clear marker of regions which become demethylated in late-stage colorectal cancer **(Figure 2c-d)**. Taken together, the analysis of DMRs suggests that 5mC and 5hmC each provide complementary information that can be used to identify dynamic methylation changes in cfDNA.

Motivated by this, we proceeded to assess whether a combination of features based on 5mC and 5hmC provided greater discriminatory power to identify stage I CRC patients from healthy controls. We built generalized linear models using the 11,691 TCGA-derived DMRs and applied a leave-one-out cross-validation (LOOCV) approach to assess model performance on stage I cfDNA samples. In LOOCV, the model is iteratively trained on all samples except one, and the left-out sample is used for testing. This process is then repeated for each sample in turn. To address variability in the learning algorithm, predictions were averaged across multiple random seeds to generate the final prediction for each test sample (see **Methods**). The classifier demonstrated an AUC of 0.54 using 5hmC only, 0.70 with 5mC only, 0.66 with modC, and 0.95 using 5mC and 5hmC (**Figure 3a**). CMS guidelines for blood-based biomarker testing for CRC detection mandate a minimum 74% sensitivity and 90% specificity. At 95% specificity, the 5mC and 5hmC model had a sensitivity of 85% (**Figure 3b**). In comparison, other publicly disclosed liquid biopsy tests reflect ongoing challenges in robust early-stage detection (**Table S1**).

**Figure 3.**
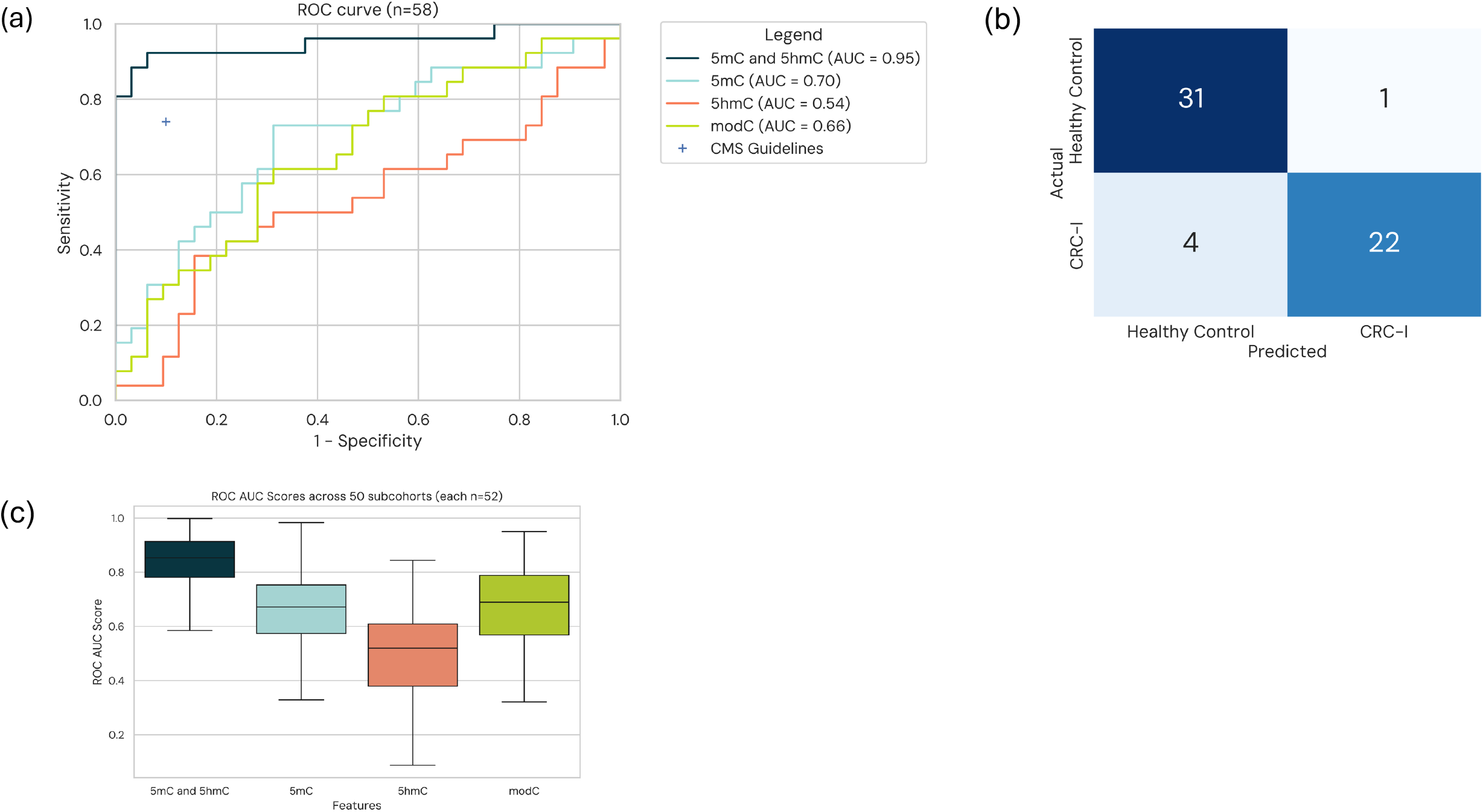
(a) Receiver operating characteristic curves for modC only, 5mC only, 5hmC only, and 5mC and 5hmC models. (b) Confusion matrix for predictions from the model using 5mC and 5hmC features with prediction threshold 0.5. (c) Box plot showing AUCs across 500 sub-cohorts for modC only, 5mC only, 5hmC only, and 5mC and 5hmC models.

To assess each model’s sensitivity to sample selection we further generated 500 sub-cohorts, each with 52 samples, by randomly excluding three CRC samples and three healthy controls per sub-cohort. For each subcohort, we repeated the LOOCV approach used on the whole cohort and computed an AUC score. Although we did observe variance in AUC across the different cohorts, models that used an independent measure of 5mC and 5hmC had consistently higher AUCs (**Figure 3c**).

We next built similar models to distinguish stage IV CRC patients from healthy controls, and to differentiate stage I from stage IV patients (**Figure S3**). In the comparison of stage IV patients with healthy controls, the highest AUC was achieved with the model using modC (AUC = 0.88), though the contribution of 5mC alone was nearly equivalent (AUC = 0.83). This suggests that 5mC provides the majority of the discriminatory signal at later stages of disease. In contrast, 5hmC contributed significantly to distinguishing between stage I and stage IV CRC, with similar AUCs for 5hmC alone (AUC = 0.95) and the combination of 5hmC and 5mC (AUC = 0.91). These results support the hypothesis that 5hmC offers additional discriminatory power in the earlier stages of cancer progression, consistent with its value in differentiating stage I patients from healthy controls. Taken together, these data suggest that models including 5mC and 5hmC as distinct biomarkers improve the performance of early-stage CRC detection.

## Discussion

In this study, we set out to evaluate whether base resolution information on 5mC and 5hmC provides additional diagnostic value, relative to 5mC alone, 5hmC alone, or modC, for stage I CRC detection using cfDNA. We observed multiple regions that transitioned from being methylated or hydroxymethylated in stage I to demethylated in stage IV CRC cfDNA. Since 5hmC is a transitional intermediate on the demethylation pathway from 5mC to C, we hypothesized that 5mC and 5hmC provided independent and additive features that could be used to improve classifier performance for early detection. Models using both 5mC and 5hmC features outperformed those based on modC, 5mC, or 5hmC alone. Notably, models used to detect stage I CRC from healthy controls frequently selected 5hmC features, highlighting the distinct contribution of 5hmC to improved classification performance (**Figure S4**).

There is potential for further studies to build on our findings and explore additional improvements. The modC DMRs were identified from microarray data from a small number of stage IV CRC tissue samples. DMR identification could be improved by assessing more tumor tissue samples, different cancer stages, and by identifying both 5mC and 5hmC DMRs. We would expect classifiers based on any of the feature sets considered here (modC, 5mC, 5hmC, and the combination of 5hmC and 5mC) to benefit from further refinement of feature selection, but we deliberately avoided this step to reduce the risk of model over-fitting and to enable clearer comparison between different models. The study employed WGS of cfDNA at a depth of 34x. Sequencing at higher depth could gather additional information, including detection of genetic variants, that may enhance both sensitivity and specificity, potentially improving overall performance.

This study leveraged the duet evoC assay to distinguish between 5mC and 5hmC, allowing for the analysis of NGS data using modC, 5mC-only, 5hmC-only, and both 5mC and 5hmC models. This flexible approach provides a powerful tool for discovery, where different indications may exhibit different levels of signal from 5mC, 5hmC, and genetics. Building on the discovery step, it is then possible to scale an assay that focuses specifically on the combination of methylation and genetics that yield the required signal (Erlitzki and Kohli, 2024).

In conclusion, distinguishing between 5mC and 5hmC enhanced sensitivity for early-stage CRC detection, showing improved performance compared to modC models. Our data supports the use of sequencing methods that distinguish 5mC and 5hmC in early detection. We feel there may be merit in applying such an approach in future studies with larger clinical cohorts, across different cancers, to more fully evaluate the potential to improve early detection performance.

## Supplementary figure legends

**Figure S1**.Histogram showing the size of hypo- and hyper-methylated DMRs from TCGA tissue data.

**Figure S2**.Scatterplots showing mean methylation differences in stage IV tissue (versus adjacent normal tissue) and mean methylation differences in stage I (upper panel) and IV (lower panel) cfDNA versus healthy control cfDNA. Each plot is annotated with the coefficient of determination R^2^.

**Figure S3**.Receiver operating characteristic curves for modC only, 5mC only, 5hmC only, and 5mC and 5hmC models for (a) stage IV CRC versus healthy control cfDNA, and (b) stage IV versus stage I CRC cfDNA.

**Figure S4**. Numbers of features selected by the LASSO classifier when using both 5mC and 5hmC as features. Each box plot represents the distribution across the 58 cross-validation splits, with the total number of features on the left, 5hmC features in the middle, and 5mC features on the right.

## Methods Cohort

This is a single center case-control retrospective cohort using cfDNA extracted from double spun plasma from consented patients undergoing routine screening for CRC by colonoscopy. Samples were obtained from National BioService LLC. Control samples were from individuals aged 45–85 years who were at average risk for CRC and had been assessed by colonoscopy, with results that showed no presence of CRC or adenomatous polyps. Cancer samples were obtained from individuals aged 45–85 years who underwent colonoscopy and were diagnosed with CRC. The following data were available for each blood sample donor: age, sex, ethnicity, and smoking status. The study was performed in accordance with the Declaration of Helsinki and was approved by the relevant independent ethics committee. Written informed consent was obtained from all donors of samples. 69 samples were selected balanced for key characteristics (age, sex and diagnosis).

### Sample preparation

Where possible, blood was sampled before colonoscopy. 10 mL blood was drawn into a K2 EDTA blood tube, placed on ice, and processed within 4 hours. Samples were centrifuged (2,000g for 10 min at room temperature). The plasma layer was transferred to a clean tube and was again centrifuged (2,000g for 10 min at room temperature) to remove any remaining cellular material. Double-spun plasma was aliquoted into tubes in volumes of at least 1 mL and then immediately frozen and stored at −80°C. cfDNA was extracted using the chemagic™ cfDNA 5k Kit (Revvity) using between 1 and 7 mL of plasma. Resulting cfDNA was quantified using a Qubit Fluorometer (Life Technologies) and quality was assessed using Agilent TapeStation and Cell-free DNA Screen Tape assay, with samples included if they had >70% for the %cfDNA quality metric on the Cell-free DNA ScreenTape assay.

### Library preparation and sequencing

The duet evoC assay was run according to manufacturer’s instructions (see Füllgrabe et al., 2023). In brief, 10ng of each cfDNA sample was end-repaired, A-tailed, and ligated to hairpin adapters. Adaptors were digested and the 3’ hydroxyl group was extended by a DNA polymerase to generate molecules with the original sample DNA strand connected to its (copied) complementary strand. Y-shaped sequencing adapters were then ligated to molecules. Methylation at 5mC was enzymatically copied across the CpG unit to the C on the copy strand using a DNA methyltransferase, whereas 5hmC was enzymatically glycosylated to prevent such a copy. 5mC on both the original and copy strands was then protected from subsequent deamination. Unmodified Cs were then deaminated to uracil, subsequently read as thymine.

After PCR, libraries were analyzed by Qubit and TapeStation and pooled at equimolar concentration. High throughput sequencing was performed on a NovaSeq 6000 using an S4 flow-cell with 2x150 reads, with 8 libraries per flow-cell, yielding an average of 1.3 billion read pairs (± 0.24 billion) per library or a mean genome coverage of 34x (SD 7.3x). Sensitivity and specificity of 5mC conversion were assessed by spike-in controls and were within expected ranges: 5mC sensitivity 96.7% ± 0.8%; 5hmC sensitivity 98.15% ± 0.5%; modC specificity 99.6% ± 0.15%. 5mC sensitivity was calculated by counting the proportion of 5mCpGs identified on a fully methylated lambda genome, 5hmC sensitivity was calculated by counting the proportion of 5hmCpGs identified on a synthetic oligonucleotide, and specificity was calculated by counting the proportion of unmodified CpGs identified on an unmethylated pUC19 genome.

Fastq files were processed using biomodal’s duet pipeline (version 1.3.0), as described in (Füllgrabe et al., 2023).

### Cohort validation

For each individual in the cohort, we assessed the agreement between the reported sex by comparing the median coverage of chromosomes X and Y. Additionally, for healthy controls and stage I CRC plasma samples, ctDNA estimates were obtained using ichorCNA (Adalsteinsson et al., 2017). All samples had estimated tumor fractions below the ichorCNA detection threshold, set at a maximum of 2.037% for healthy controls and 4.6% for stage I CRC patients. This threshold was not applied for stage IV CRC plasma samples.

### Feature generation

DNA methylation data of paired tumor and normal samples of patients with stage IV CRC was downloaded from TCGA (https://portal.gdc.cancer.gov/, January 2024). These included methylation data of 9 paired tumor-normal samples using Illumina Infinium Human Methylation 450K BeadChip (Illumina 450K array). Methylation probes without any beta values in any sample were filtered out. DMRs between stage IV and normal samples were identified using DMRcate (Peters et al, 2021). Significant DMRs with FDR < 0.05 were lifted over from hg19 to GRCh38 genomic coordinates. Epigenetic data (modC, 5mC, and 5hmC fractions) was summarized across regions identified as significantly differentially methylated in paired stage IV CRC tissue samples. Average modC, 5mC, and 5hmC fractions for each region were obtained by dividing the sum of the counts of each cytosine modification by the count of all cytosines across all CpGs in the reads covering each region of interest.

### Model building

Model building was carried out using the R package Glmnet (Friedman et al., 2010; Tay et al, 2023). Briefly, given the high dimensionality of the dataset (∼11k features x 58 samples) a regularization step was performed to decrease the number of features used to build each model. Feature selection was carried out using a lasso logistic regression model. The regularization strength lambda was chosen by cross-validation on the training set, with the number of folds set to 10. The best lambda value was chosen as *lambda*.*min*, unless no features were retained, in which case lambda was chosen as the minimum lambda returning a model with at least 1 feature.

For evaluation of the different feature sets, a LOOCV approach was used. In this method, the model is trained on all but one sample in the dataset and tested on the remaining sample. This process is repeated for each sample in turn, meaning that each sample serves as a test set once. To address variability in the training process, predictions were averaged across 25 random seeds. The predicted probabilities (corresponding to the probability a sample was stage I CRC according to the model) for each sample were used to generate the ROC curves and AUC scores. To address the reproducibility of the results to different starting cohorts, 500 different sub-cohorts were generated by removing a random subset of 6 samples (3 with stage I CRC and 3 healthy controls) from the main cohort and evaluating each feature set.

### DMR calling

Differential modification (i.e. 5mC, 5hmC, or modC) calling in cfDNA data was carried out by aggregating counts of modified and unmodified cytosines across all CpG contexts within the regions defined by the analysis of stage IV TCGA samples.

A logistic regression model was employed to analyze the aggregated data. The model is structured as follows assuming *N* samples separated into two groups. For each sample *i*, the proportion of modified bases in the region *p*_*i*_ is modelled via the logistic regression model:

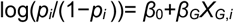

In this equation, *β*_*0*_ is the intercept and *β*_*G*_*X*_*G*,*i*_ represents the group term (e.g., stage IV vs. control, where *X*_*G*,*i*_ would be 0 if sample *i* belongs to the control group, and 1 if it belongs to the stage IV group).

For each region of interest, we perform an independent statistical test to evaluate the null hypothesis H_0_: *β*_*1*_ = 0. If we reject the null hypothesis, it indicates that the log-odds (and consequently, the modification proportions) differ significantly between the treatment and control groups. In this case, we classify the region as a differentially 5mC or 5hmC modified region. Conversely, if we fail to reject the null hypothesis, it suggests that there is no statistically significant difference in modification levels between the two groups for that particular region. DMR calls were corrected for multiple testing using Benjamini-Hochberg, resulting q-values were called based on a predetermined p-value threshold. DMRs were called for both 5mC and 5hmC separately.

**Table S1.** Sensitivity and specificity for CRC stage I detection. N is the number of samples assessed. ND is not disclosed. Grail refers to the Circulating Cell-free Genome Atlas study (NCT02889978; Klein et al, 2021), Freenome to the PREEMPT study (NCT04369053), and Guardant to the ECLIPSE study (Chung et al, 2024).

| Reference | Stage I |  |  | Stage IV |  |  |
| --- | --- | --- | --- | --- | --- | --- |
|  | Specificity | Sensitivity | N | Specificity | Sensitivity | N |
| Grail | 99.5% | 43.3% | 30 | 99.5% | 95.3% | 64 |
| Freenome | 91.5% | 57.1% | ND | ND | 100% | ND |
| Guardant | 89.9% | 55% | 22 | 89.9% | 100% | 10 |

## Acknowledgements

The results published here are in whole or part based upon data generated by the TCGA Research Network: https://www.cancer.gov/tcga.

## Notes

### Competing Interest Statement

S.B. is a founder, adviser and shareholder of biomodal Ltd. All the other authors are current or former employees and hold share options. Patents covering this work have been filed by biomodal Ltd.

