## Supplementary figures and images for "5-methylcytosine and 5-hydroxymethylcytosine are synergistic biomarkers for early detection of colorectal cancer"

### Supplementary Figures

Supplementary Figure 1

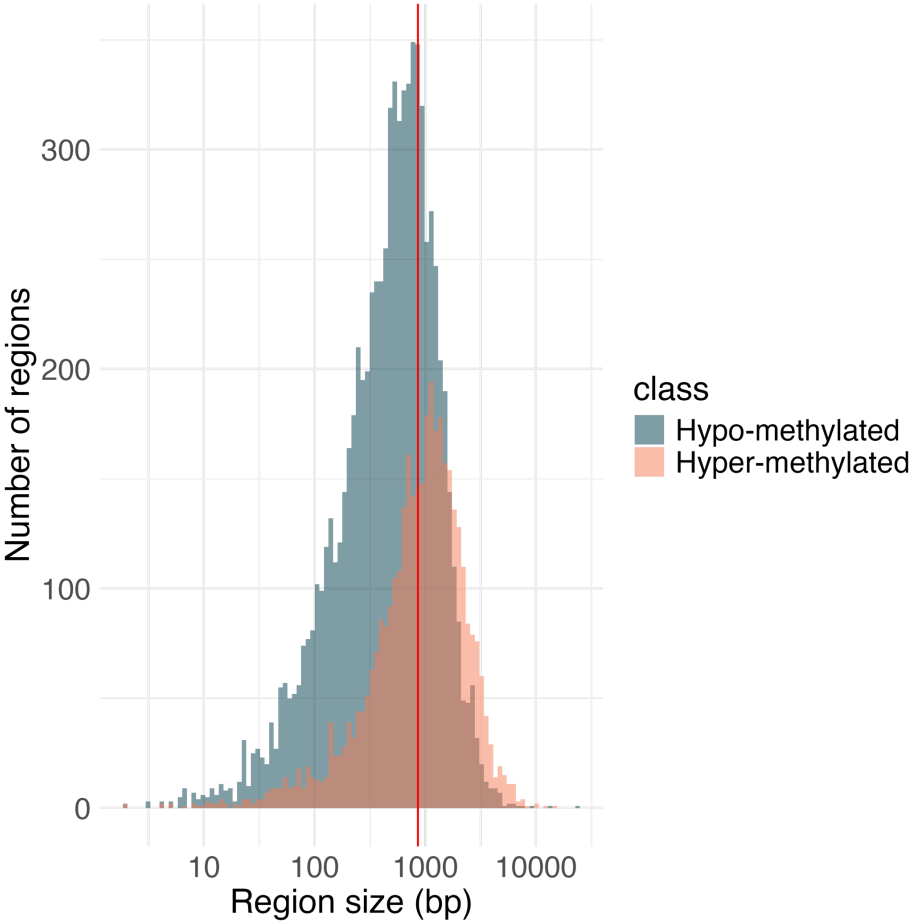

#### Supplementary Figure 2

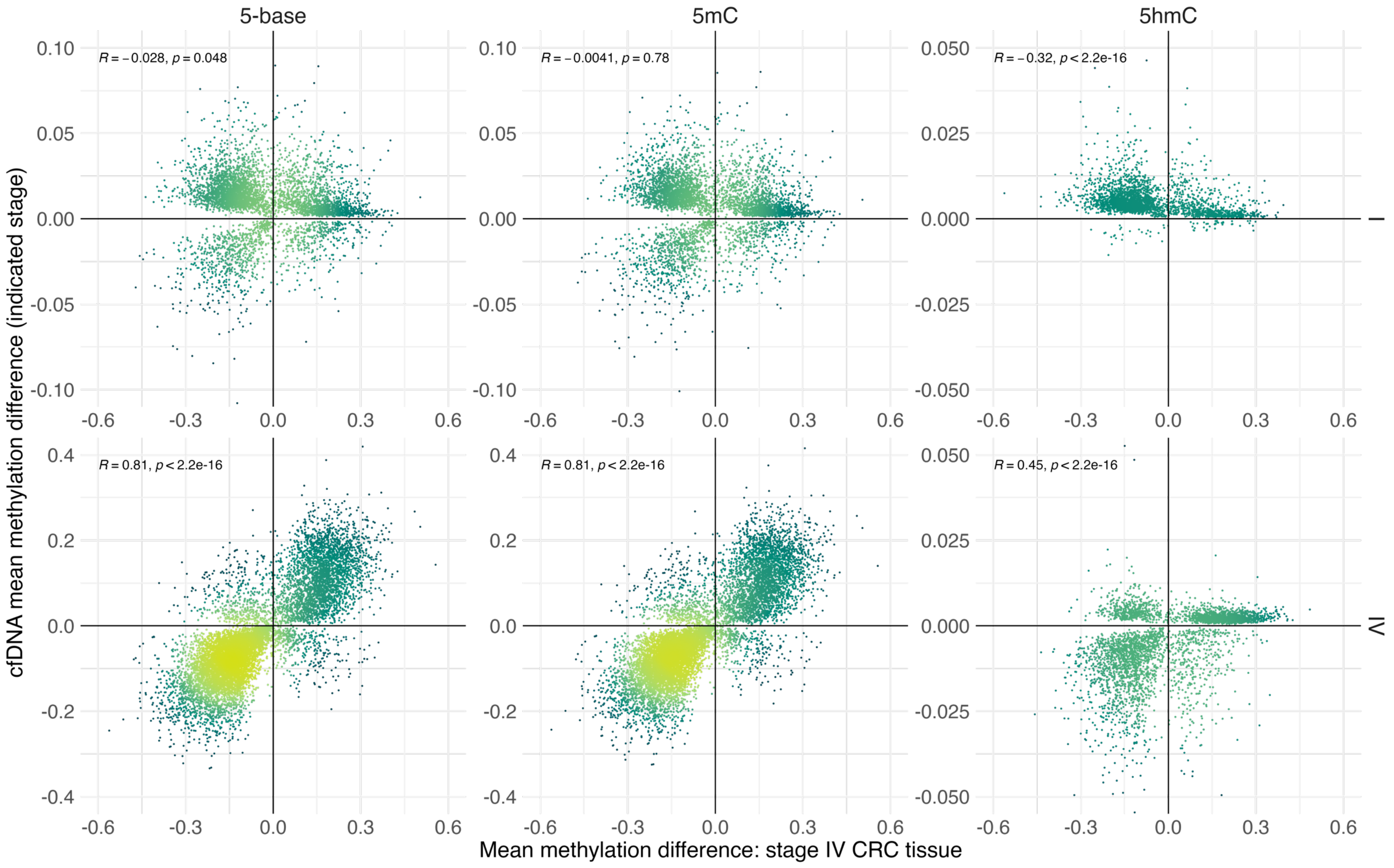

Supplementary Figure 3

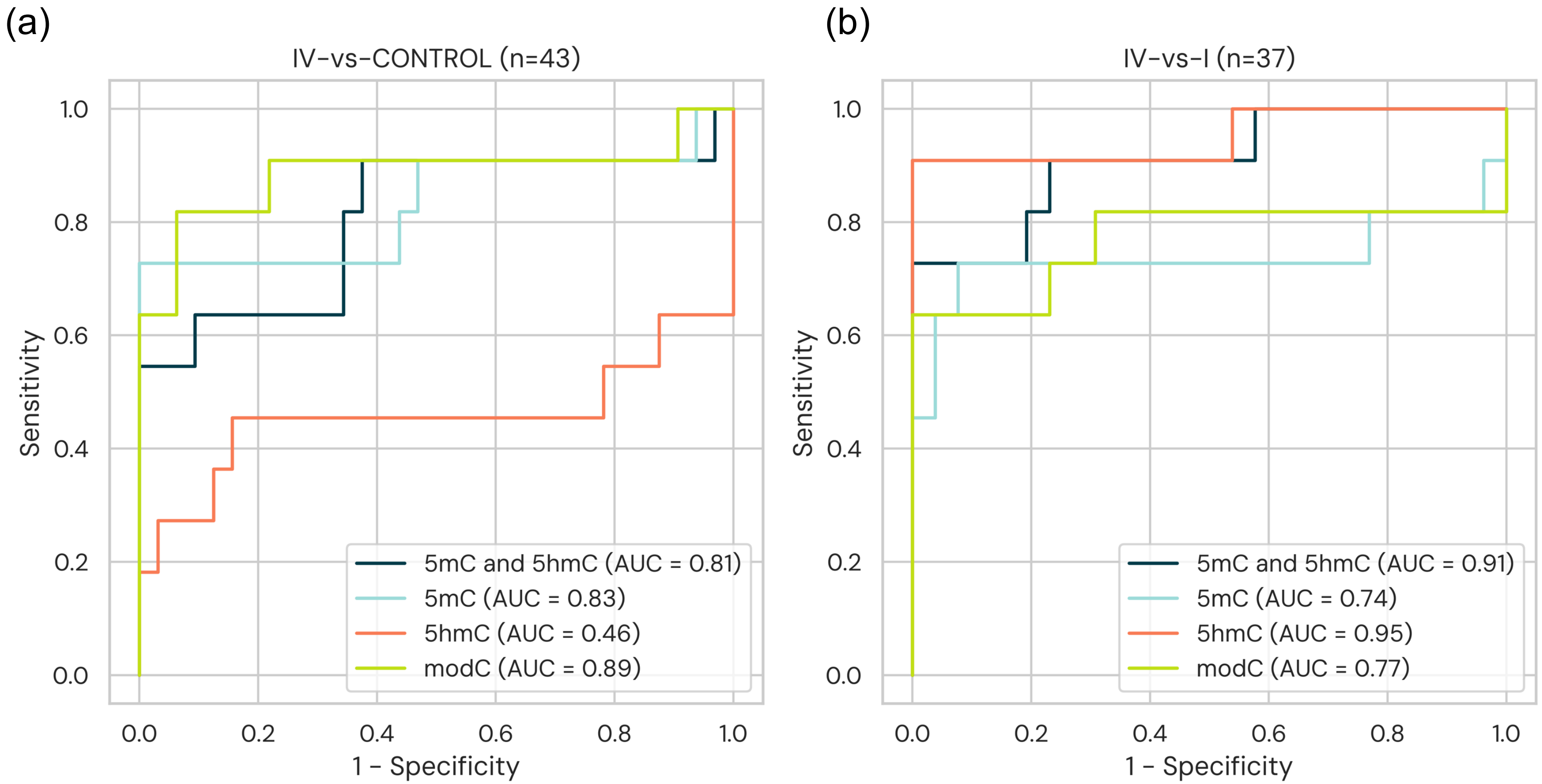

Supplementary Figure 4

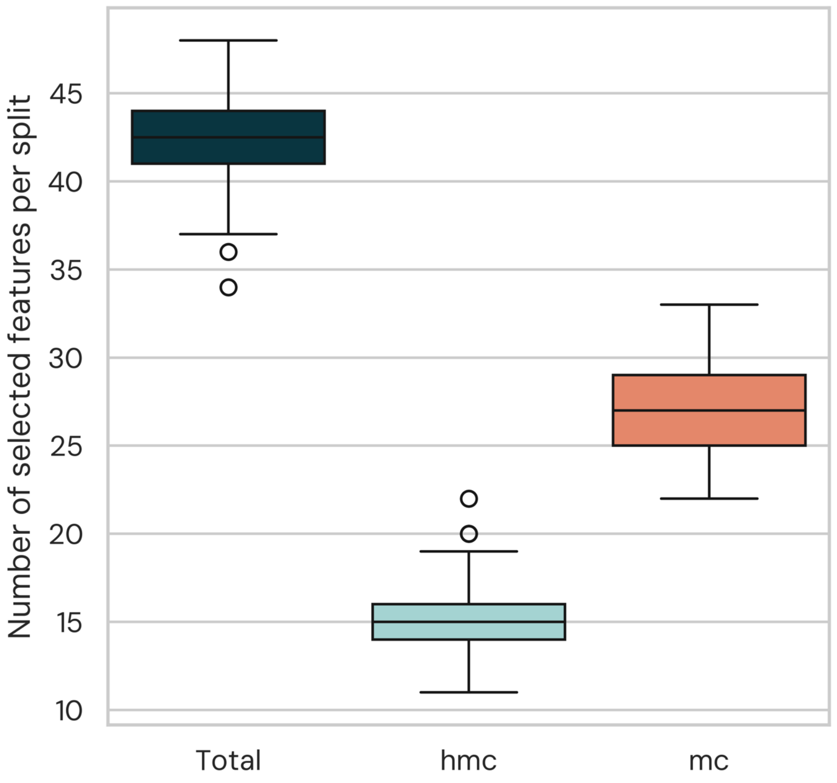
